# Nose-to-ground distance drives the evolution of mammalian odorant receptor repertoire size

**DOI:** 10.64898/2025.12.23.695377

**Authors:** Joël Tuberosa, Fandilla-Marie Furlan, Alan Carleton, Ivan Rodriguez

## Abstract

The largest protein-coding gene family in mammals encodes the repertoire of chemosensors responsible for detecting odorants, namely odorant receptors (ORs). OR gene numbers vary widely across species, ranging from a few hundred in some primates to over two thousand in elephants. Here, we report that proximity to the ground, a rich source of olfactory cues, has broadly influenced the evolutionary expansion and contraction of OR mammalian repertoires. Analyzing 158 terrestrial mammalian species, we find a strong inverse correlation between nose-to-ground distance and the number of functional OR genes. In contrast, no significant association was found between nose-to-ground distance and the size of other chemoreceptor families, including vomeronasal and bitter-taste receptors. As a case study, we examined OR repertoire divergence between giraffe and okapi lineages, revealing that differential rates of gene duplication and pseudogenization resulted in giraffes possessing 25% fewer ORs than okapis. Together, our findings suggest that the chemical complexity of the olfactory environment has shaped the evolution of OR repertoires, providing new insights into gene family size and diversification in mammals.

## Introduction

Mammals rely on their olfactory and gustatory systems to perceive and interpret the chemical diversity of their environment. The ability to detect a wide array of compounds relies on hundreds of genes coding for chemosensory G protein-coupled receptors (GPCRs), each with distinct ligand affinities. These receptors have originated from multiple lineages within the GPCR phylogeny, integrating into chemosensory systems and expanding into numerous paralogs (*1*, *2*). The four largest chemoreceptor gene families include bitter taste receptors (T2Rs) (*3–5*), type 1 and type 2 vomeronasal receptors (V1Rs and V2Rs, mostly devoted to pheromone detection; (*6*, *7*)) and odorant receptors (ORs; (*8*)), which represent the most expansive gene family among the four (*9*).

OR genes are expressed in the nose, in the main olfactory epithelium, where each sensory neuron expresses a single OR gene from a single allele (*10*, *11*). Since multiple receptors can respond to the same compound and a single receptor can recognize multiple ligands, odor perception is thought to result from a unique combinatorial code of receptor-ligand interactions (*12*). Neurons expressing the same OR converge in the olfactory bulb and form homotypic sensory channels (*13*, *14*). There, the signals are integrated (*15–17*) with further processing occurring in the cortex to generate distinct odor perceptions (*18*).

The number of OR genes varies remarkably across species, ranging from approximately three hundred to two thousand. This results from fluctuating rates of birth-and-death evolution (*19*, *20*). Alongside this process, mutations in OR sequences may alter ligand affinities (tuning) and thus shift detection spectra (*21*, *22*).

Changes in habitat, diel activity or diet may have channeled the OR repertoire evolution, in terms of size (*20*, *23–26*) and tuning (*27*).

A reduction in selective pressure to maintain sensory function is usually associated with loss of the corresponding sensors. For instance, marine mammals have experienced significant reduction in their OR repertoire size (*25*, *28*, *29*). Similarly, among primates, a dietary shift from frugivory to folivory has been linked to an accelerated decline in functional OR genes (*30*). A comparable trend is observed in the visual system, where transitions to nocturnality or dim environments correlate with opsin gene degeneration (*31–34*). As a corollary, the expansion of sensor repertoires would result from adaptive selection. Consequently, a nocturnal lifestyle is thought to drive olfactory adaptation towards large OR repertoires in mammals (*27*, *31*, *35*) as well as in birds (*36*, *37*). Beyond the number of genes, selective pressure can also affect receptor tuning. In primates, the shift from nocturnal to diurnal activity has been associated with OR tuning specificity loss, alongside changes in opsin sensitivity (*27*).

However, these pressures that possibly affected OR repertoire sizes only explain the OR evolution of very specific mammalian phylogenetic branches.

The concept of sensory drive (*38*) proposes that species exposed to complex chemical environments evolve more sophisticated chemosensory systems (*39*). In terrestrial environments, chemical complexity is highest at ground level, where organic compounds released by plants, fungi, bacteria, and animals accumulate (*40–43*). Odorants deposited on the ground form horizontal gradients that macrosmatic mammals such as rodents and canines can track (*44–46*), whereas plumes rising from the ground are rapidly disrupted by atmospheric turbulence (*47*).

Based on the hypothesis that species with limited exposure to ground-level odors may experience relaxed selective pressures on the number and complexity of their olfactory sensors, we investigated a potential link between the distance separating the nostrils to the ground and the chemosensory gene repertoire size across 158 species covering the whole phylogenetic mammalian tree. We found a significant and general association for OR repertoires, underscoring the critical role of environmental chemical complexity in shaping the evolution of the largest gene family in mammals.

## Results

### Diversity of OR gene repertoire sizes in mammals and variability of nose to ground distances

We first constituted a dataset of 158 species representing all orders of terricolous mammals for which quality genomic data were available (n=18 orders; Supplementary Table 1). To identify OR coding sequences in their corresponding genome assemblies, we designed a pipeline to search and validate homologous OR coding sequences using TBLASTN, hidden Markov models, and critical protein motives (Supplementary Figure 1). We identified 197,543 OR coding sequences, of which 137,659 were classified as functional, with the remaining considered pseudogenes. Consistent with previous reports, the number of genes coding for functional ORs varied substantially across terrestrial mammals, ranging from 295 genes in the grivet (*Chlorocebus aethiops*, 1.5% of all protein-coding genes) to 2,348 genes in the Asian elephant (*Elephas maximus*, 12% of all protein-coding genes).

To account for phylogenetic dependence, we estimated rates of molecular evolution across lineages. We identified 1,900 nuclear ultra-conserved elements loci in our genome set, totaling 1,576,939 aligned sites. Lineage genetic divergence was estimated by performing maximum likelihood estimation of branch lengths in a consensus mammal phylogeny. Using the resulting species tree, we identified groups of closely related species (separated by less than 0.01 substitution per site) and sampled a single representative taxon from each group. The resulting species set included 96 species (Figure 1A,B). This set is referred to as the main species set, with 250 alternative species sets generated to establish confidence intervals for statistical evaluations.

**Figure 1.**
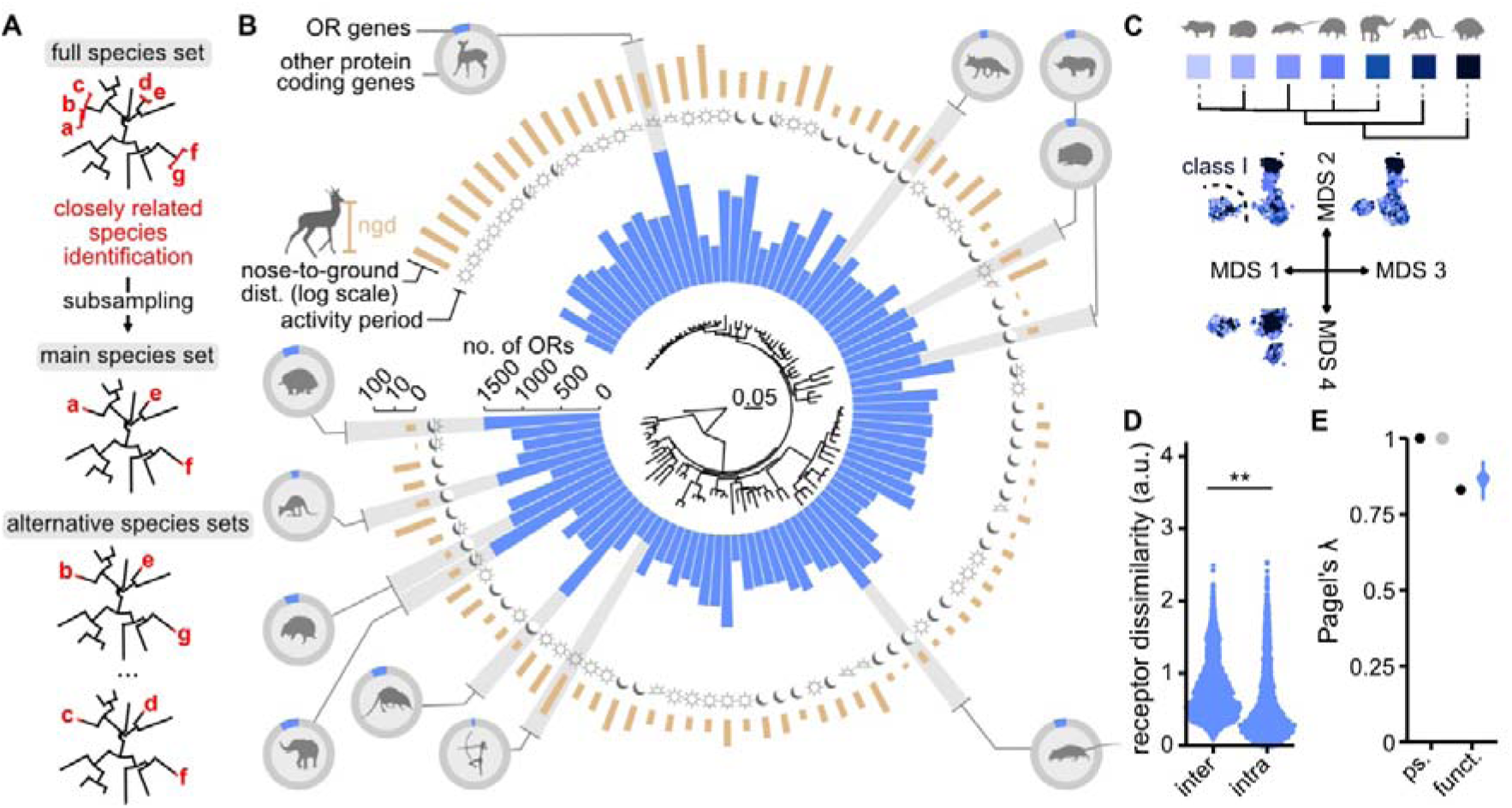
OR gene repertoire sizes in mammals. (**A**) Generation of the main and alternative species sets by subsampling single representative species within closely related species clades. (**B**) In the center, a phylogeny of 96 representative mammal species. The topology was obtained from the Timetree website (*50*), the branch lengths were estimated by maximum likelihood using ultraconserved elements (1,576,939 sites) to estimate overall genetic divergence between species. Tree scale: substitution/site. Species data of functional OR gene repertoire sizes (no. of ORs, blue bars), activity periods (sun symbol: diurnal, moon crescent: nocturnal, half sun: crepuscular, moon and sun: unspecific) and nose-to-ground distances (ngd, in logarithmic scale), are aligned the phylogeny. Ten species are represented in silhouettes, with the proportions of genes coding for functional ORs in all protein coding genes (from top left, clockwise: *Muntiacus reevesi*, *Vulpes vulpes*, *Ceratotherium simum*, *Erinaceus europaeus*, *Rattus norvegicus*, *Homo sapiens*, *Rhynchocyon petersi*, *Loxodonta africana*, *Tolypeutes matacus*, *Macropus fuliginosus, Tachyglossus aculeatus*). (**C**) Distribution of OR proteins from seven species after multidimensional scaling of their pairwise dissimilarities into three dimensions. Species affiliations are shown in shades of blue next to their phylogeny (scale: substitution/site). (**D**) Density distribution of dissimilarity scores between the closest pairs of receptors, belonging either to two different species (interspecies) or to the same species (intraspecies). The Cohen’s D is displayed on the top. ***: p-value < 1e-16. (**E**) Phylogenetic signal (*λ*) values carried by the number of functional OR (blue) or OR pseudogene (grey), calculated over 250 alternative species sets, with the main set *λ* shown in black. Other values are represented as quantile ranges. Dot: median; large segments: interquartile range; thin segments: 95% confidence interval.

To assess OR repertoire content divergence, we compared 8,985 OR sequences from seven representative species (Supplementary Figure 2A). Pairwise differences between these sequences were calculated based on amino-acid chemical properties of the region spanning from transmembrane domain one to seven, which is believed to include the ligand-binding pocket. Multidimensional scaling revealed that mammalian ORs cluster primarily by phylogenetic classes rather than species of origin (Figure 1C), consistent with their phylogenetic distribution (*48*). Nevertheless, some of the short-beaked echidna (*Tachyglossus aculeatus*) ORs appeared to form a cluster diverging from the rest of the repertoires, potentially indicating an elevated proliferation rate in this OR gene lineage.

To quantify species-specific OR identity, for each receptor we identified its most similar counterpart, either within the same species or in a different species. The average difference in amino-acid similarities between these categories, was estimated using Cohen’s D measure (mean difference scaled by the pooled values standard deviation), yielding a value of 0.63 (Figure 1D), with ORs from the same species showing greater similarity compared to cross-species comparisons.

While most OR identities are shared across mammals, the number of functional OR genes exhibits substantial variability. To determine the strength of the relationship between OR number fluctuation and phylogenetic distance, we calculated Pagel’s λ – a phylogenetic signal measure (*49*) – across the 250 alternative species sets. The resulting values ranged between 0.79 and 0.92 (Figure 1E), meaning that variance in OR number deviates from that expected in a Brownian motion model (where λ=1). Conversely, similar estimation for OR pseudogenes consistently showed a strong phylogenetic signal (λ=1; Figure 1E), suggesting that functional OR gene number fluctuations result from heterogeneous proliferation rates decoupled from pseudogene accumulation, potentially reflecting differential selective pressures across lineages.

For each species, exposure to ground-level odors was estimated by the measure of the distance between the tip of the nose and the ground at rest (ngd, see Material and methods). The ngd ranged from 0 (mole species) to 4.21 m (*Giraffa camelopardalis*). Species in the dataset were predominantly diurnal (n=68) or nocturnal (n=60), with the remaining classified as crepuscular (n=15) or with unspecified activity patterns (n=15).

### Nose-to-ground distance predicts the number of functional OR genes

To test the association between nose-to-ground distance (ngd) and the number of functional OR genes, we performed a phylogenetic generalized least square regression (pGLS) using the logarithm of ngd. The regression was highly significant in our main species set (likelihood ratio test vs. null model, p-value = 0.00168; Figure 2A,B). Notably, this association was not observed for the number of OR pseudogenes (Figure 2C).

**Figure 2.**
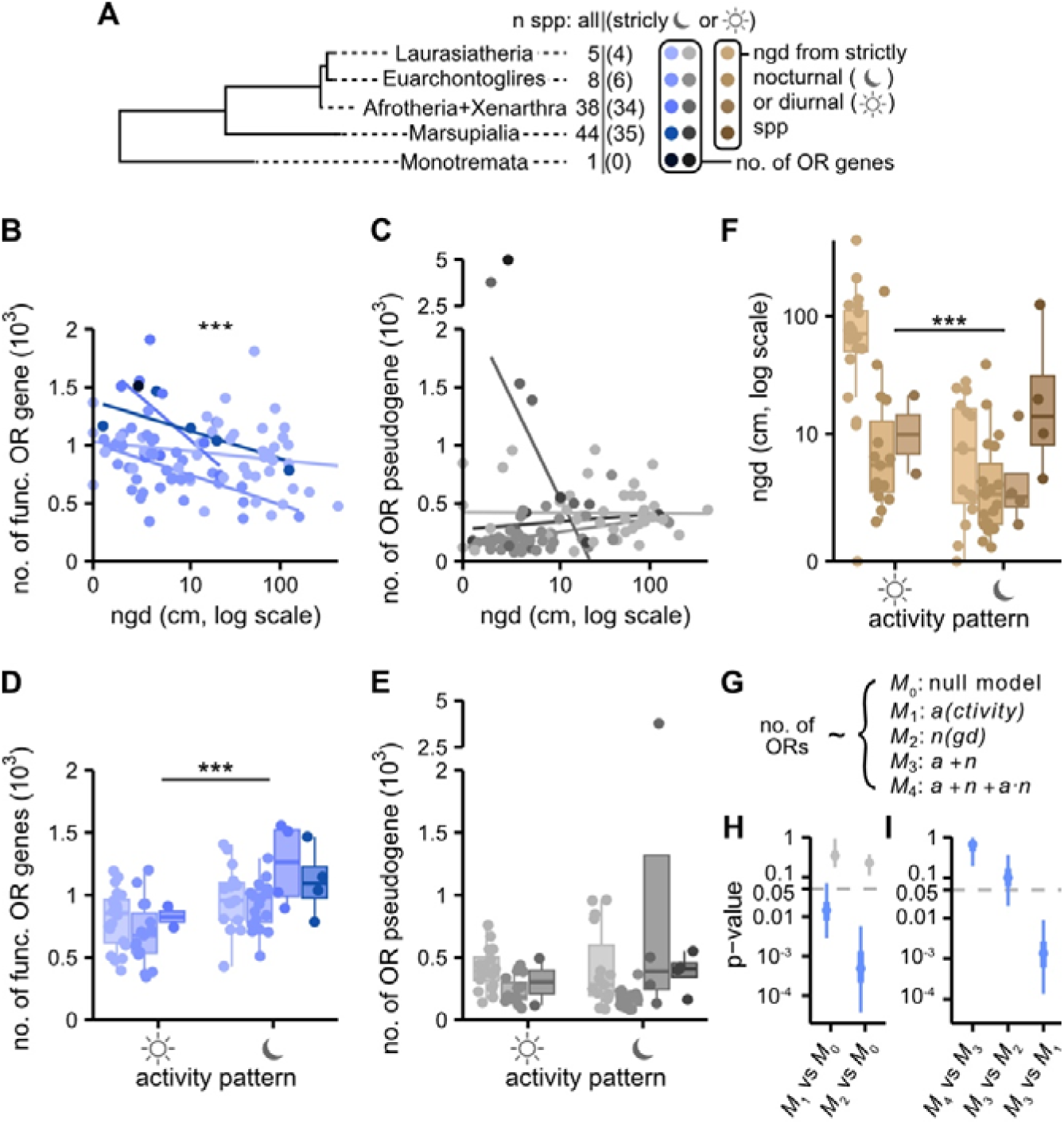
Nose-to-ground distance better explains OR repertoire size variation than the activity period. (**A**) Species data in the next plots were colored according to their membership in the different phylogenetic groups illustrated here. Statistics in **B**, **C** and **H** (*M*_1_ vs *M*_0_) were computed with all species, regardless of their activity period. Statistics in **D**, **E**, **F**, **H** (*M*_2_ vs *M*_0_) and **I** were computed using only species recorded with strict nocturnal or diurnal activity periods. (**B**) Functional OR gene number as a function of the nose-to-ground distance (ngd); ***: pGLS p-value = 0.00168 with n=96 species (main set). Linear regressions (without phylogenetic correction) are shown for four different phylogenetic groups (all groups but the Monotremata, *Tachyglossus aculeatus*). (**C**) OR pseudogene number as a function of the ngd; n.s: pGLS p-value = 0.249 with n=96 species (main set). (**D**) Functional OR gene number as a function of the activity period (strictly nocturnal or strictly diurnal); ***: pGLS p-value = 0.00438 with n=79 (subset of the main set, see **A**). (**E**) OR pseudogene number as a function of the activity period (strictly nocturnal or strictly diurnal). n.s: pGLS p-value = 0.152 with n=79 (subset of the main set). (**F**) Ngd as a function of the activity period (nocturnal or diurnal). n.s: pGLS p-value = 0.00427 with n=79 (subset of the main set). The same grouping applies as for panel **D** and **E**. (**G**) Key of the regression models compared in **H** and **I**. The response variable is always the number of OR genes (functional or pseudogenic). Each of the non null model (*M*_1-4_) are described by their explanatory variables (*a*: activity period; *n*: nose-to-ground distance). (**H**) Ranges of pGLS p-values obtained with the alternative sets for the two alternative models (*M*_1_, with n=96 species, and *M*_2_, with n=79 species). (**I**) Ranges of pGLS p-values obtained with the alternative sets by comparing nested models, using only species that are strictly nocturnal or diurnal (n=79).

Our finding was apparently not congruent with the main current explanation for modulating OR gene repertoire sizes in mammals: adaptation to dark environments (*27*, *31*, *35*). We first evaluated whether nocturnal lifestyle is indeed associated with a higher number of functional ORs. We refined our dataset to include only strictly diurnal or nocturnal species, excluding crepuscular and unspecified activity categories. The pGLS regression of the category means confirmed that nocturnal species possess a significantly higher number of ORs (p-value = 0.00438, Figure 2D), a trend that was absent in pseudogene counts (Figure 2E).

However, subsequent analyses revealed a significant correlation between ngd and activity period (p-value=0.00427; Figure 2F), suggesting that one dimension may simply reflect the other. To address potential confounding effects and disambiguate the predictive power of these variables, we compared multiple statistical models: A full model including ngd, activity period, and their interaction term, a model without the interaction term and individual models for ngd and activity period (Figure 2G). To evaluate the consistency of the p-value, we replicated these analyses across the 250 alternative datasets. When examined independently, both predictors demonstrated significant associations with functional OR gene numbers across most datasets. The ngd showed consistent significance, while the activity period was non-significant in 6% of alternative species sets (Figure 2H). These variables were never found to be associated with pseudogene counts (Figure 2H). The full model was not significantly better at predicting OR numbers than the model without the interaction term (two-variable model) and could therefore be rejected (Figure 2I). Critically, in 80.8% of the cases, the two-variable model performed equivalently to the ngd-only model, suggesting that the activity period does not increase the model’s predictive power. Conversely, the two-variable model was systematically superior to the activity-only model (Figure 2I). These results suggest that the predictive power of activity period on functional OR gene numbers is largely mediated through its correlation with ngd. Ultimately, nose-to-ground distance emerges as the more robust predictor of functional OR gene numbers in mammals.

### Repertoire sizes in other large chemosensor gene families

To explore whether similar pressures would influence other large chemosensor repertoires, we analyzed the distribution of the number of genes coding for vomeronasal receptors type 1 (V1Rs) and type 2 (V2Rs), and bitter taste receptors (T2Rs). We extracted the corresponding genes from the genomes of the 158 mammalian species, using a similar pipeline than that used for the OR gene identification (Supplementary Figure 1). Like the OR repertoire, the number of functional genes coding for these receptors (5277 V1Rs, 3261 V2Rs and 1897 T2Rs in total) exhibited substantial variation across mammalian species (Figure 3A).

**Figure 3.**
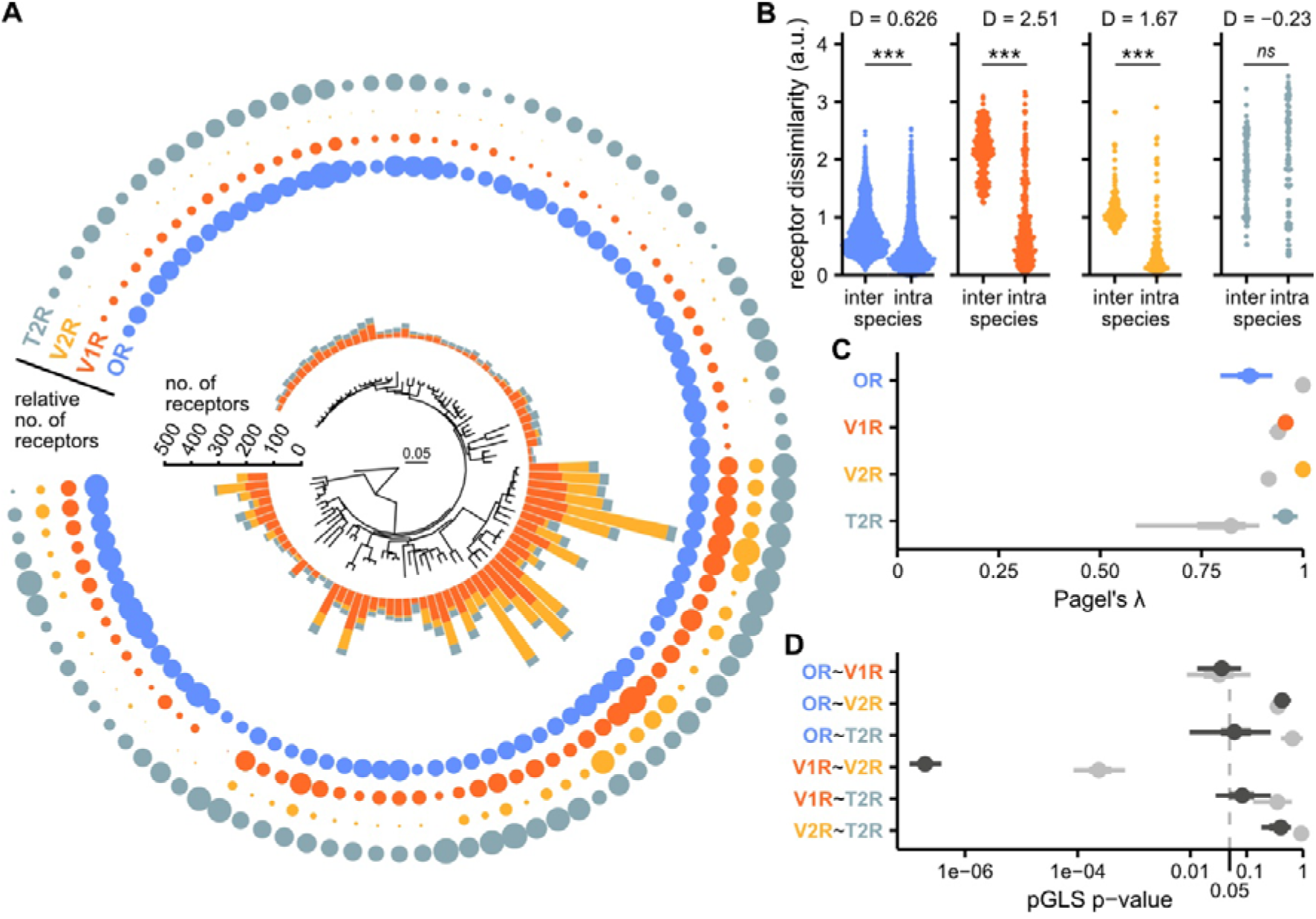
Differential rates of evolution across olfactory and gustatory receptor repertoires. (**A**) Repertoire sizes of odorant receptors (OR), pheromonal receptors (V1R and V2R) and bitter taste receptors (T2R) of 96 representative mammals (the main species set, see Figure 1D). Bars: absolute number of functional receptors (without the ORs), circles: relative numbers, by receptor family. (**B**) Distribution of dissimilarity levels between the closest pairs of receptors from a given family in different (inter-) or in the same (intra-) species. The repertoires of seven representative mammals were used for this analysis (same as in Figure 1B; n=8985 ORs, 396 V1Rs, 200 V2Rs, 106 T2Rs; ***: p-value < 1e-16, n.s: p-value > 0.05). (**C**) Phylogenetic signal of the repertoire size of each family (colored dots and line ranges: functional genes, light grey: pseudogenes). (**D**) Ranges of the p-value of pGLS regressions computed with each pair of repertoires (dark grey: functional genes, light grey: pseudogenes).

Rodent genomes contain high number of vomeronasal receptor genes – for example, *Mus musculus* has 227 V1Rs, and the Mearns’s grasshopper mouse (*Onychomys arenicola*) has 389 V2Rs. This pattern is also observed in marsupials – for instance, the gray short-tailed opossum (*Monodelphis domestica*) possesses 88 V1Rs and 106 V2Rs. Conversely, several mammalian orders (Artiodactyla, Carnivora, Macroscelidea, and Primates) are nearly devoid of V2Rs. Likewise, bitter taste receptor (T2R) functional gene counts exhibited remarkable variation, ranging from a single gene in the short-beaked echidna (*Tachyglossus aculeatus*) to 43 genes in the prairie vole (*Microtus ochrogaster*).

To assess divergence in receptor identities across repertoires of different mammalian lineages, we performed the same analyses as described earlier for the OR genes. By pooling 8,985 ORs, 396 V1Rs, 200 V2Rs, and 106 T2Rs (Supplementary Figure 2A), we compared the pairwise similarity between receptors of the same species versus receptors from two different species. Notably, V1Rs and V2Rs displayed a higher degree of species-specific identities than ORs (Cohen’s D = 2.51 and 1.67, respectively; Figure 3B). In contrast, T2Rs exhibited a distinct pattern, with indistinguishable levels of dissimilarity between intra- and inter-species receptor pairs, indicative of high ortholog conservation and low gene duplication rates.

Multidimensional scaling and hierarchical clustering of pairwise dissimilarity matrices (Supplementary Figure 2B) further confirmed these observations. The analysis revealed a higher degree of species-specific clustering in V1Rs and V2Rs compared to ORs and T2Rs (Supplementary Figure 2C-J), potentially resulting from accelerated birth-and-death rates in the two pheromonal receptor gene families (*48*) and their prominent involvement in intra-specific interactions, accelerating shifts in the repertoire tuning (*51–53*).

Chemosensory receptor affinity is linked to amino acid sequence identity. Consequently, sequence diversity is associated to some degree with detection spectrum breadth. To quantify the relationship between receptor gene numbers and sequence diversity, we constructed a minimum spanning tree of amino acid sequences from each individual species repertoire and extrapolated the level of receptor diversity from the total tree length (Supplementary Figure 3A). Receptor diversity showed a strong correlation with the number of genes in the repertoire (Pearson correlation test: p-value < 1e-16; Supplementary Figure 2L-O), suggesting that larger gene repertoires expand the potential detection spectrum through increased sequence variation.

Phylogenetic signal analysis using Pagel’s λ revealed deeper clade dependencies for V1R, V2R, and T2R gene numbers compared to ORs. The median λ values were 0.95 for V1Rs, 1.0 for V2Rs, and 0.95 for T2Rs, significantly higher than those observed for olfactory receptors (Figure 3C).

Receptors pertaining to the same sensory organ can be expected to be exposed to similar challenges. Phylogenetic generalized least squares (pGLS) regression across alternative datasets consistently demonstrated that functional V1R and V2R gene counts were significantly correlated, with a median p-value of 1.94e-7 (adjusted by Holm’s method). This correlation extended to their pseudogene numbers (median adjusted p-value = 2.39e-4). Other receptor family pairs exhibited more variable correlation patterns across datasets.

### The nose-to-ground distance is weakly – if at all – associated with the number of V1R, V2R, or T2R genes

Exposure to an increasingly complex chemical environment might be expected to exert similar selective pressures on different modalities of chemosensory perception, including odorant, pheromonal, and gustatory perception. However, the systems supporting these functions differ in anatomy, ligand specificity and access (in particular, ligand volatility), and perceptual outcomes. To assess whether, like for ORs, proximity to the ground influences the evolution of pheromonal and gustatory receptor repertoires, we regressed the number of V1R, V2R, and T2R genes against nose-to-ground distance (Figure 4A-D). Additionally, we gathered from publicly available resources several trait variables that could affect the evolution of these different gene repertoires, including activity period, diet, territoriality, and sociality (see Material and methods), and evaluated their potential association with chemosensory repertoire sizes. After correction for multiple testing, only the associations between the number of functional ORs and either nose-to-ground distance or activity period remained statistically significant (Figure 4A-D).

**Figure 4.**
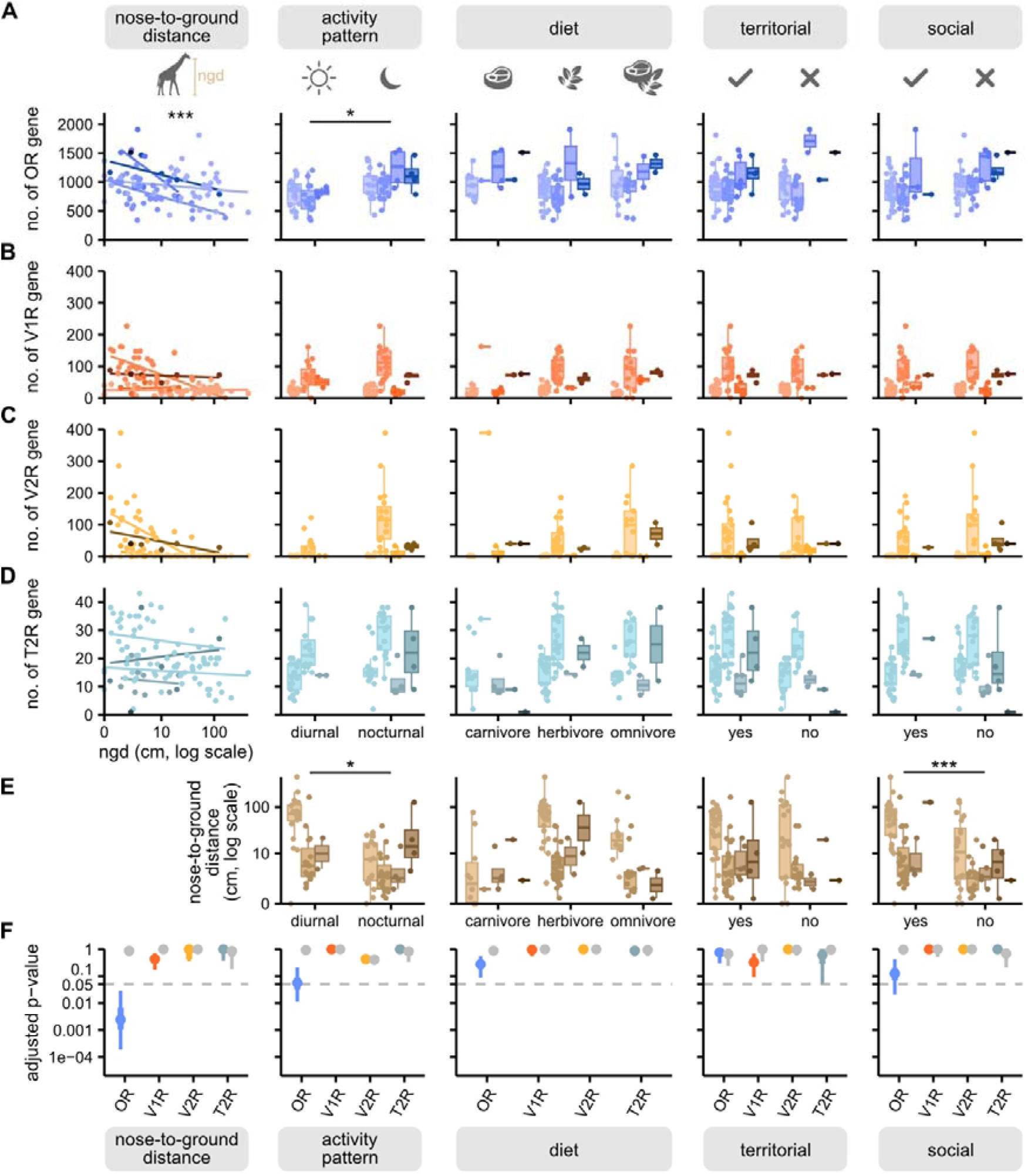
Nose-to-ground distance best explains variance in the number of functional OR genes (but no other chemosensory genes) relative to other trait variables. (**A**-**D**) Number of functional genes (**A**: ORs, **B**: V1Rs, **C**: V2Rs, **D**: T2Rs) with regards to different traits characterizing each species of the main species set. From left to right: the animal’s nose-to-ground distance (in logarithmic values, n=96), the animal’s activity period (strictly nocturnal or diurnal, n=79), the animal’s diet (eats animals, eats plants, or eats both, n=96), whether the animal is territorial or not (n=96), and whether the animal is social or solitary (n=96). Significance of statistical tests (pGLS for all variables except diet, phylogenetic one-way ANOVA for the diet): **: p-value < 0.01, *: p-value < 0.05, n.s: p-value > 0.05. P-values were corrected for multiple comparison with the Holm method. (**E**) Comparison of trait variables with the nose-to-ground distance. The same statistical tests were conducted than for the number of functional chemosensory genes (n=76 for the activity period, and 96 for the other traits). From **A** to **E**, boxplots and regression lines are drawn for phylogenetic groups shown in Figure 2A. (**F**) P-value ranges of the regression of the number of functional genes by the different traits, using the alternative datasets.

While the linear relationship between nose-to-ground distance and the number of OR genes appeared to hold, we considered that the relationship could be valid within a limited range of distances for other receptor families. Notably, species with very short nose-to-ground distances seem to exhibit higher numbers of pheromone receptor genes (Figure 4B,C). To further investigate this, we compared the mean number of chemoreceptor genes between species classified as having either short or long nose-to-ground distances. By incrementally adjusting the distance threshold, we found that the mean number of ORs became significantly different when the threshold was set at ≥4 cm (Supplementary Figure 4A). Among pheromone receptors, no significant association was found for V1Rs (Supplementary Figure 4B); however, the mean number of V2R genes was significantly higher in species with a nose-to-ground distance <2.5 cm (Supplementary Figure C). These species (n = 24) are primarily rodents (n = 13), followed by laurasiatherian insectivores (n = 5) and six additional species from different mammalian orders. No association was found for functional T2R genes, although the mean number of T2R pseudogenes was significantly higher in species with a nose-to-ground distance >52 cm (Supplementary Figure 4D).

To identify potential confounding effects, we tested the association between nose-to-ground distance and each of the other potentially explanatory trait variables (Figure 4E). In addition to the previously observed association between nose-to-ground distance and activity period (Figure 2F), we found that sociality was also significantly associated with nose-to-ground distance. However, sociality only had a marginally significant effect on the variance in OR gene number (adjusted p-value = 0.054), which we therefore did not consider further.

To assess the robustness of these findings, we repeated all tests using 250 alternative species datasets. In these analyses, the association between nose-to-ground distance and the number of functional OR genes remained highly significant (99.6% of adjusted p-values < 0.05; Figure 4F). By contrast, the association between activity period and OR gene number was significant in only 43.8% of cases, while all other tested associations were significant in at most 14.7% of cases (e.g., OR number and sociality, Figure 4F).

Following the findings of Padilla-Morales et al. (*54*), who reported a positive association between body size and the number of functional OR genes, we re-evaluated this hypothesis within our analytical framework. This association was significant in 41% of our alternative datasets (Supplementary Figure 5). However, unlike nose-to-ground distance or activity period, this association was also significant for OR pseudogenes in 54% of alternative datasets (Supplementary Figure 5). No association was found for other gene families, nor between sexual size dimorphism and nose-to-ground distance. These results suggest that the observed association remains uncertain and that the link with pseudogenes may reflect an influence on gene duplication rates rather than selective pressures on coding sequence conservation.

In summary, our findings strongly support nose-to-ground distance as a key driver of OR repertoire dynamics, a variable that does not appear to affect the repertoire of other chemoreceptor superfamilies, to the possible exception of V2Rs.

### Repertoire size difference between the giraffe and the okapi is the result of differential rates of gene duplication and pseudogenization

To examine a case of OR repertoire divergence between two closely related species that differ in height but share similar diets and activity periods, we compared the OR repertoires of the okapi (*Okapia jonhstoni*, nose-to-ground distance ∼140 cm), a rainforest browser, and the giraffe (*Giraffa camelopardalis*, nose-to-ground distance ∼420 cm), a savanna browser. These species diverged approximately 16 million years ago from an ancestor that was roughly the same size as the okapi (*55*, *56*). The giraffe possesses 920 OR genes, including 476 functional genes (48% pseudogenes), while the okapi has 986 OR genes, with 626 coding for functional receptors (37% pseudogenes). To investigate the underlying gene birth-and-death dynamics driving this divergence, we reconstructed the phylogeny of giraffe and okapi OR genes (Figure 5A; Supplementary Figure 5). We then identified all orthologous gene lineages forming fully supported sister clades (n = 395, bootstrap scores >90%) and inferred ancestral states to retain only the gene lineages for which the most recent common ancestor was a functional gene (n = 347).

**Figure 5.**
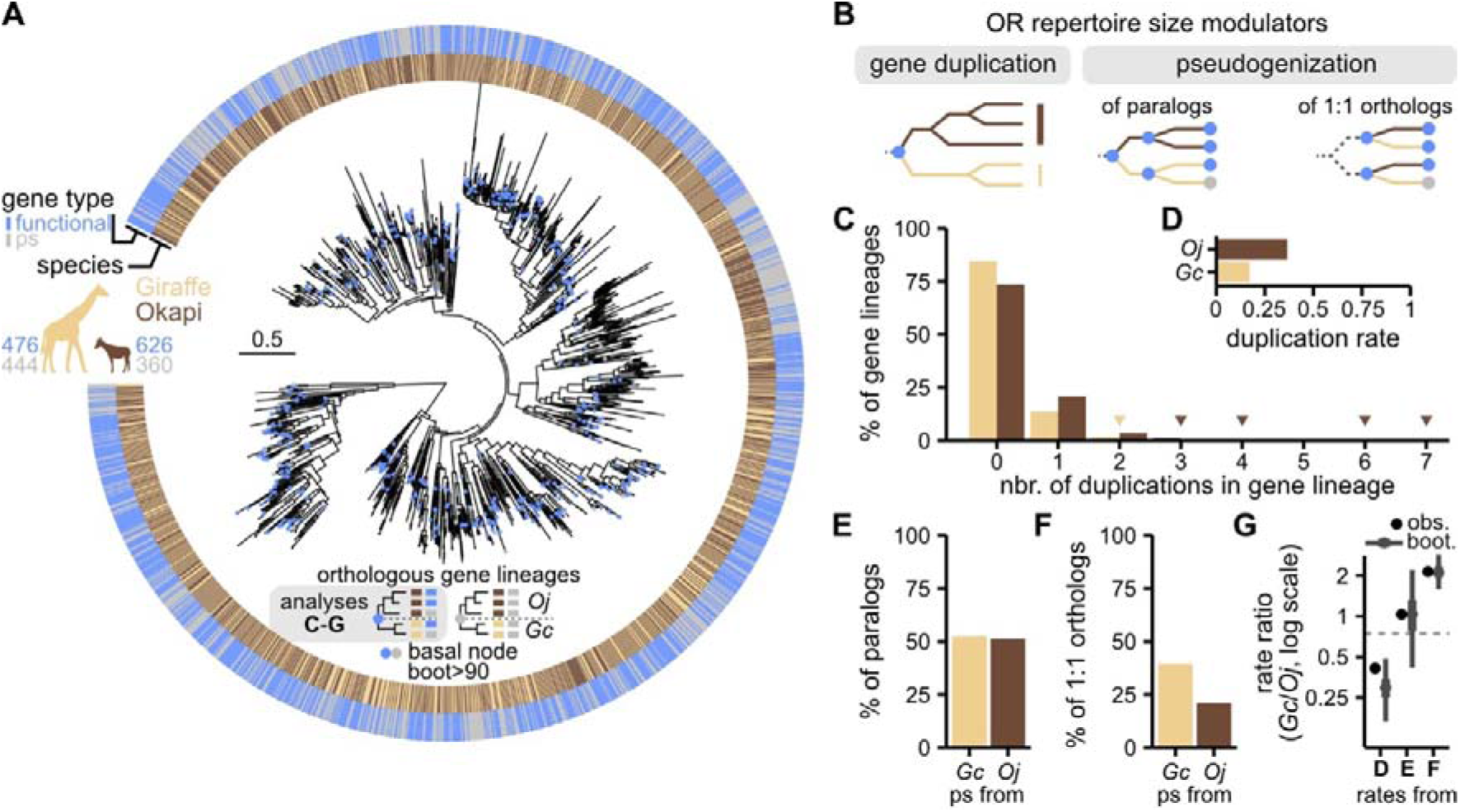
The difference in OR repertoire sizes between the giraffe and the okapi results from differential gene birth-and-death rates. (**A**) Maximum likelihood phylogeny of odorant receptor functional genes and pseudogenes of the giraffe (*Giraffa camelopardalis*) and the okapi (*Okapia johnstoni*), n=2006 genes. The inner ring indicates species affiliation with colored tiles. Dark brown: okapi; gold: giraffe. The outer ring indicates the gene type. Blue: functional; grey: pseudogene (ps). The numbers next to the species silhouettes correspond to gene counts of each gene type and for each species. Scale in amino-acid substitution per site. Dots on the phylogram designate nodes (bootstrap > 90%) corresponding to the coalescence of two orthologous gene lineages. Blue node: inferred functional OR ancestor. Grey node: inferred OR pseudogene ancestor. Clades originating from functional OR ancestors constitute the data analyzed in the next panels. (**B**) Two processes modulate the number of functional OR genes over time: gene duplication and pseudogenization. Pseudogenization rate was measured for two different gene categories: genes derived from duplication events (paralogs) and gene lineage that did not duplicate (1:1 orthologs). (**C**) Percentage of gene lineages (y-axis) that have undergone a specific number of duplication events (x-axis). (**D**) Percentage of genes derived from duplication in the giraffe (*Gc*) and in the okapi (*Oj*) lineage. (**E**) Percentage of duplication-derived genes that became pseudogenes in each lineage. (**F**) Percentage of one-to-one (1:1) orthologs that became pseudogenes in each lineage. (**G**) Rate ratios derived from the observed values in D (duplication rate), E and F (pseudogenization rates). Black dot: observed value. Grey range: 95% of bootstrapped values. The dashed line shows the expected divergence due to the differential genetic divergence of the okapi and the giraffe lineage.

The difference in functional gene numbers between the okapi and the giraffe could be attributed to variations in gene duplication frequencies and/or pseudogenization rates (Figure 5B). Our data indicate that a minority of gene lineages underwent duplication: the proportion of duplicated gene lineages was 15.4% in the giraffe and 26.7% in the okapi (Figure 5C). However, 63.3% of the okapi’s OR genes (including 39.2% pseudogenes) arose from duplications following divergence from the giraffe, compared to only 28.7% in the giraffe (including 49.3% pseudogenes; Figure 5D). Among paralogs, the proportion of gene duplications in which one of the resulting copies became a pseudogene reached 52.4% and 51.3% in the giraffe and okapi genomes, respectively (Figure 5E). For one-to-one orthologs, the giraffe exhibited a higher pseudogenization rate (39.5%) than the okapi (21%; Figure 5F).

To evaluate whether gene duplications or pseudogenizations events accumulated at different rates between the giraffe and the okapi lineages, we compared the rate ratios of these events to the molecular evolutionary rate ratio inferred with the UCE-calibrated mammalian phylogeny (MRCA to *Giraffa camelopardalis*=5.44e-3; MRCA to *Okapia johnstoni*=7.33e-3 substitution/site; giraffe/okapi=0.742). We found that the gene duplication rate ratio between the giraffe and the okapi (0.412) was lower than the expected molecular evolutionary ratio (Figure 5G). To assess the robustness of this signal against sampling bias, we performed 250 bootstrap iterations on the orthologous gene lineage set. The bootstrap value range never overlapped with the expected molecular evolutionary ratio, indicating that the rate of gene duplication was significantly lower in the giraffe lineage than in the okapi lineage. Then, we applied a similar test to evaluate pseudogenization rates in both duplicated and non-duplicated genes. We could not reject that pseudogenization rates among duplication-derived genes were different between the two lineages, as the expected rate ratio fell within the bootstrap value range (79.6% of which was above the expected ratio; Figure 5G). However, the giraffe lineage exhibited a significantly higher pseudogenization rate among one-to-one orthologs, with all bootstrap values exceeding the expected rate ratio (Figure 5G).

Taken together, these results suggest that the divergence in OR repertoires between the giraffe and the okapi was driven by significant differences in both selective pressures maintaining functional ORs and gene duplication rates.

## Discussion

In this study, we investigated the evolutionary drivers of the genes encoding ORs, the largest gene family in the mammalian genome. We asked whether the size of this exceptionally large repertoire of sensory genes may have evolved under a sensory drive. To test this, we focused on terrestrial mammals, in which the strength of the sensory drive was represented by nose-to-ground distance – the distance between the probed gaseous volume and the richest, most concentrated source of volatile compounds. We selected 158 terrestrial mammalian genomes, from which we extracted the respective numbers and identities of functional genes and pseudogenes for the four major chemosensory gene families pertaining to different chemosensory modalities: ORs (odorant detection), V1Rs, V2Rs (pheromone detection) and T2Rs (bitter-taste compound detection).

Comparing repertoire size evolution across chemoreceptor families, we found that the OR repertoire size exhibits less phylogenetic signal than the other chemoreceptor families. This is indicative of a higher degree of species-specific birth-and-death evolution dynamics. On the other hand, when comparing sequence identities within and in between species, we observed that the V1R and the V2R repertoires display a higher degree of species-specific identities compared to OR or T2R repertoires, consistent with their higher rate of gene duplication (*48*). As vomeronasal receptors modulate intraspecific interactions, it is likely that sexual selection has driven shifts in V1R and V2R repertoire tuning (*51–53*). Thus, whereas the number of vomeronasal receptors may retain deeper phylogenetic dependency than ORs, the identities composing these repertoires evolved much faster. Finally, T2R repertoires show a higher degree of shared identity across mammals than species-specific identity, a high conservation of orthologous sequences suggesting a limited and shared identity of T2R ligands across species.

Concurring with the sensory drive hypothesis (*39*) and based on our comparison of repertoire sizes with nose-to-ground distance as well as other putatively relevant ecological factors, we found that this distance is negatively correlated with the number of functional OR genes. From this perspective, species with greater nose-to-ground distances would tend to be exposed to a less complex olfactory environment and thus to be more prone to accelerated pseudogenization of OR genes compared to species living closer to the ground, leading to repertoire shrinkage. This would constitute a confounding effect to the previously proposed sensory trade-offs between vision and olfaction during the evolution of arboreal mammals as well as in giraffids, for whom it is proposed that selective pressure shifted from the olfactory to the visual system due to the increasing predominance of vision (*57*, *58*).

Earlier reports suggest that nocturnal lifestyle may drive olfactory adaptation in mammals towards larger OR repertoires – despite the fact that trade-off between vision and olfaction is the least pronounced across mammals compared to other modality pairs (*57*). This potentially compensatory link with activity during light phases is reminiscent of the hypothesis that the evolution of a highly developed olfactory system in the ancestor of extant mammals was associated with a shift towards nocturnality (*31*), as well as the observation that nocturnal or cave-dwelling species often possess an exceptional sense of smell (*37*, *59*). Our findings, however, seem to contradict a primary role of activity pattern in shaping OR repertoire size. Although we did observe an association between diel activity and repertoire size, this association was nonetheless weaker than that observed with nose-to-ground distance. We propose a straightforward explanation for this diel activity signal, suggesting a direct – and thus confounding – link between activity patterns and olfactory exposure: nocturnal mammals tend to be short in stature (Figure 2F), and their noses are therefore closer to the ground.

Our study was designed to identify ecological drivers of OR repertoire size evolution acting broadly across terricolous mammals. By construction, our statistical framework and sampling scheme did not allow the detection of clade-specific effects and may have overlook dynamics taking places at narrower phylogenetic scales such as dietary shifts in primates (*30*) or altitude-related niche adaptation in Boreoeutherian placentals (*26*).

The causal relationship between nose-to-ground distance and OR repertoire size could be considered reciprocal. If a larger and more diverse OR repertoire enhances the ability to parse complex olfactory environments, it is plausible that OR proliferation would open up ecological opportunities, by making ground-level foraging, nesting or shelter-seeking more accessible. In this context, extensive OR repertoires may in some cases have driven morphological adaptations, such as elongated snouts, downward-oriented nostrils, or reduced stature.

With the exception of the association between nose-to-ground distance and OR repertoire size, all associations between repertoire sizes from the four tested families (OR, V1R, V2R and T2R) and the five trait variables were lost after correction for multiple testing and phylogenetic dependency. However, for V2Rs, a significant association for the nose-to-ground distance emerged when comparing animals with very short nose-to-ground distances (less than 2.5 cm) to the others. This result may reflect the fact that V2Rs bind molecules that are less volatile than V1R or OR ligands – many V2R ligands are known to be peptides – and thus require contact or near-contact with the source for detection.

To dissect the dynamics of gene birth-and-death evolution in a specific case of OR repertoire divergence, we compared the OR genes of the okapi and its cousin, the giraffe, which is three times taller. Their most recent common ancestor lived around 16 million years ago. We found that the giraffe possesses 37% fewer functional OR genes than the okapi. After reconstructing the phylogenies of orthologous OR gene lineages, we observed a higher rate of gene duplication in the okapi compared to the giraffe. Seven gene lineages underwent more than three duplications in the okapi lineages, versus none in the giraffe. This apparent heterogeneity may reflect bursts of gene duplication at specific loci, or the selection of advantageous locus multiplication. To assess pseudogenization rates, we measured pseudogene frequencies in both duplicated and non-duplicated genes. In both categories, the giraffe exhibited a higher pseudogene frequency than the okapi. Unlike duplication rates, these pseudogenization rate ratios corresponded to the median of bootstrap values, suggesting a relatively homogeneous process. Our results indicate that the pressure to maintain functional ORs has been more relaxed in the giraffe lineage compared to the okapi, while gene duplication has occurred more frequently in the okapi lineage. Taken together, these results suggest that the sizes of their OR repertoire more likely reflect differential selective pressure – acting on the whole repertoire – rather than genetic drift. In light of our previous findings, the height of these species may have influenced their repertoire size evolution given different levels of sensory drive.

In conclusion, our analysis assessed the role of sensory drive in shaping the evolution of the OR gene family across terrestrial mammals. It suggests that sensory environments complexity, more than sensory trade-off, has translated into selective pressure affecting birth-and-death rates of the largest mammalian gene family, thus impacting the genome on a broad scale.

## Supporting information

UCE-calibrated species phylogeny, full species set (n spp=158).

UCE-calibrated species phylogenies, 250 alternative species sets (n spp=96).

Coordinates of functional and pseudogenic receptor coding sequences.

Giraffe and okapi OR phylogeny.

## Data and code availability

All custom scripts, Hidden Markov models, coordinates of functional and pseudogenic coding sequence alignments, and phylogenies generated in this study will be available upon publication.

## Acknowledgements

We thank Dr Jose Manuel Nunes for expert technical assistance in statistical design. We thank all members of I.R and A.C laboratories for helpful discussions. We thank Ivan Ishchenko from https://animalia.bio/ for kindly providing source data table from his website. We gratefully acknowledge Joel Sartore for the invaluable resource that is the Photo Ark. This research was supported by the Swiss National Science Foundation (grants 310030_215572 and 501100001711-219531 to I.R., and 310030_219531 to A.C).

## Author contributions

Ivan Rodriguez and Alan Carleton, conceptualization, funding acquisition, supervision, writing – original draft, review & editing; Joël Tuberosa and Fandilla-Marie Furlan, investigation, project administration, formal analysis, visualization, writing – original draft, methodology.

## Material and methods

### Genome assembly selection

To constitute a representative set of mammalian genomes, we listed terricolous mammalian representative genome assemblies using the NCBI dataset CLI tool (*60*). The resulting genome assembly set underwent completeness quality control using Busco (*61*) using either published scores or running the quality control ourselves, using Busco v5.8.0 with the MetaEuk pipeline. Assemblies with a completeness score under 80% were filtered out. This resulted in a set of 158 species (Supplementary Tables 1-2).

### Nose-to-ground distance and other species traits

Information about dimensions, habitat, activity period, diet, sociality and territoriality of each species was retrieved from the University of Michigan’s online database of animal diversity of (https://animaldiversity.org/) or when information was missing, from the Encyclopedia of Life (https://eol.org) or Animalia (https://animalia.bio/). Trait lists and descriptions were manually reviewed and for each trait, animals were sorted into one of the corresponding categories, as listed in Supplementary Table 3. To measure the nose-to-ground distance, for each species, we selected one image of a representative adult individual at rest that we collected from various open-license sources and the Photo Ark gallery (https://www.joelsartore.com/gallery/the-photo-ark/; all images with dimension lines: Supplementary Data 1). The nose-to-ground distance in cm was then extrapolated given the average height or length of the species. For sociality and territoriality, cases of sexual dimorphism were classified as ‘social’ or ‘territorial’, respectively. Trait data (including the nose-to-ground distance) for each species are found in Supplementary Table 4.

### Species phylogeny and genetic distances

Ultra conserved elements were identified and aligned using the software PHYLUCE version 1.7.3 (*62*) and following the procedure described under the Tutorial III: Harvesting UCE Loci From Genomes and the Tutorial I: UCE Phylogenomics, Finding UCE loci. Briefly, we used a 5k probe set optimized for tetrapods (*63*) to probe the genome assemblies listed in Supplementary Table 2. All identified loci were then aligned, the resulting alignments were trimmed and cleaned using the default parameters. Alignments containing less than 95% of the taxa were filtered out. Molecular divergence between species was then estimated by maximum likelihood. For this, we downloaded the time-calibrated species phylogeny from the TimeTree website (*50*) in February 2024. Then, we used IQ-TREE version 2.0.7 (*64*) to perform a maximum likelihood optimization of the branch lengths from the UCE alignment partitioned by locus. Substitutions rates were estimated using the model GTR+I+G, whose parameters were optimized for each alignment partition. The resulting tree was used to calculate the covariance matrix of the generalized least squares (GLS) regressions, that is to perform phylogenetic GLS regressions (*65*).

### Alternative species sets

To generate alternative species sets and their corresponding phylogeny, the following subsampling procedure was implemented. In the three containing all species, we identified nodes including species distant from less than 0.01 substitution per site from each other. A closely related species group was formed with the taxa found after each of these nodes. These nodes were pruned, and alternative trees were formed by sampling one species in each of the closely related species groups and regrafting the corresponding terminal branch to its original parent node. Cophenetic distances and tree manipulations were calculated and performed in R using functions of the ape package (*66*). Phylogeny of the comprehensive species set is available as Supplementary Data 2. Alternative sets phylogenies are available as Supplementary Data 3. Species subsets featured in the different analyses are detailed in Supplementary Table 1.

### Coding sequence identification

To constitute a probe set for homology search, we retrieved protein sequences of OR, V1R, V2R and T2R of four mammalian species, the mouse (*Mus musculus*), the cow (*Bos taurus*), the African elephant (*Loxodonta africana*), and the nine-banded armadillo (*Dasypus novemcinctus*) from Ensembl (*67*), using InterPro annotation (OR: IPR000725, V1R: IPR004072, V2R: IPR004073, T2R: IPR007960). In V2R, the sequence coding for the N-terminus region preceding the first transmembrane domain spans several exons and is highly variable. Thus, we trimmed the probe sequences from the conserved motif CxxC to retain only the last part of the protein, which lays on a single exon. Subsequent sequence identification and curation steps only consider sequences aligned to this region. To limit redundant searches, we excluded each protein having a homolog with more than 80% of sequence identity. The resulting probe sets was then used for TBLASTN searches in all genomes. Protein integrity was assessed by detecting seven transmembrane domains using TMHMM 2.0. The probe sets were then aligned with MAFFT G-INS-i algorithm and trimmed before the most conserved start methionine and after the last conserved positions (30% conservation of the most common amino acid; and for V2R, starting from the CxxC motif). Sequences of the probe sets were used as TBLASTN queries to identify homologous sequences in our genome set. For this procedure, we set the best hit filtering options - best_score_edge 0.3 and -best_hit_overhang 0.3 and kept only hits with an E-value lower than 1e-20. Then hits distant of less than 30 bp or overlapping were merged using two different merging procedures. One consists in merging hits in the same reading frames (respective to the genome sequence) and extracting the longest open reading frame overlapping with this range and therefore identifying putatively functional coding sequences (which, for V2R, was considered from the conserved motif CxxC). The other consists in merging hits regardless of the reading frame, while compensating with artificial insertions to recover the original reading frame, and therefore to pseudogenic coding sequences. Regarding the putatively functional coding sequences, a series of verifications was performed to assess whether they would code for a functional receptor of the targeted family. First, a hidden Markov model was designed from the each of the probe set alignments with hmmbuild from the hmmer package (*68*) version 3.3.2. A statistic of the score was obtained by scanning the probe set sequences with their respective model using hmmscan. For each model, a threshold score was calculated as the third of the average score of this statistic. Putative coding sequences of each receptor family were scanned with their corresponding model and ruled out if they displayed a score below the threshold. Transmembrane domain integrity was checked in the remaining sequences by using TMHMM (*69*) version 2.0c and only sequences with 6 to 8 detected transmembrane domains were retained. Finally, the remaining sequences were manually reviewed and discarded if they displayed a deletion larger than 10 amino acids and if they exhibited degeneration in homologous sites of highly conserved motifs (Supplementary Figure 1). As for the pseudogenic coding sequences, we assessed their identity using the hidden Markov models as described for the putatively functional coding sequences. To retain only the pseudogenic sequences, previously identified functional coding sequences were subtracted from these sets. Numbers of functional or pseudogenic receptors for each species are found in Supplementary Table 4. The coordinates of the corresponding sequences are found in Supplementary Data 4.

### Chemoreceptor pairwise dissimilarity

Receptor protein sequences were trimmed to retain the sequence from the first to the last transmembrane domain based on transmembrane domain prediction by DeepTMHMM (*70*). Pair of receptors were aligned with needle from the EMBOSS package (*71*). Dissimilarity between each homologous amino-acids were scored according to the Miyata amino acid replacement matrix (*72*). For each non-ambiguous amino-acid aligned with a gap or an ambiguous amino-acid, the dissimilarity was calculated as the mean dissimilarity score for divergent pairs including that amino acid. For each ambiguous amino acid aligned with a gap or another ambiguous amino acid, the score was calculated as the average of all possible dissimilarity scores. The sum of the dissimilarity scores collected alongside the alignment of a given pair of receptors constituted the dissimilarity score for these receptors.

Metric multidimensional scaling of the pairwise distance matrix into a 3-dimensional space was computed in R with the function cmdscale, with option k=3.

To compare the dissimilarity between receptors within a given species’ repertoire with the dissimilarity of receptors coming from two different species’ repertoires, we first collected, for each receptor, the closest receptor in the same species and the closest receptor in a different species. This resulted in two distributions per chemoreceptor gene family, that we compared using the Cohen’s D value, which was calculated with the cohen.d function from the package effsize (*73*).

To quantify the level of species-specific receptors, we performed a Ward’s hierarchical clustering (*74*) from the pairwise dissimilarity matrix of the seven species pooled repertoires. For this, we used the hclust function in R, with the Ward.D2 algorithm. Next, we extracted all species-specific clusters by identifying all nodes of the resulting tree leading to receptors from a single species. The size of the cluster was established by counting the number of receptors they contained.

To estimate the chemoreceptor diversity within repertoires of individual species, pairwise dissimilarity matrices were obtained for each analysed repertoire using the same method as described above. From these matrices, minimum spanning trees were computed with the minimum_spanning_tree function of the SciPy package (*75*), which uses the Kruskall algorithm. Total tree lengths were obtained by summing all the edge lengths of a given graph.

### Phylogenetic signal and regressions

Phylogenetic generalized least square regression (pGLS) where performed with the function phylolm from the R package of the same name (*76*), in R 4.4.2 (*77*). The function was always provided with the UCE-calibrated phylogeny and ran with the option model=’lambda’, which indicates to perform a maximum likelihood estimation of Pagel’s λ, scale internal phylogeny branches accordingly, and calculate the covariance matrix from the resulting phylogenetic distances.

To evaluate the range of Pagel’s λ values obtained across alternative dataset, gene counts were regressed on a null model using the phylolm function, and λ values were extracted from the function results. For statistical testing of univariate models using either the ngd, the activity period, the sociality or the territoriality as the explanatory variable, likelihood ratio tests were performed by comparing the likelihood of the univariate model with that of the null model. Likewise, the full model and the multivariate model implying ngd and activity period were compared to their nested counterparts to evaluate the significance of their predictive power difference.

For the evaluation of the link between the repertoire size and the diet, or between the ngd and the diet, we performed a phylogenetic ANOVA (*78*) in R, with the phylANOVA function of the phytools package (*79*) and using the same species phylogeny as for the pGLS tests.

All corrections for multiple testing were performed using the Holm–Bonferroni method.

### Giraffe and Okapi repertoire phylogeny

Protein sequences of functional ORs and translations of CDSs classified as pseudogenes of the two species were grouped by class (I or II). Class I and Class II sequences were aligned separately with Clustal Omega (*80*) using option --full and --full-iter, and the two alignments were subsequently aligned as profiles with the same program. The alignment was trimmed from the first and the last position showing 40% of identity, leaving sequences covering the seven transmembrane domains with an overhang of about 15 amino acids on each end.

The OR phylogeny was inferred using IQ-TREE version 2.0.7 (*64*). Three amino-acid substitutions models were tested with the ModelFinder algorithm: WAG, JTT and LG including either four discrete gamma rate categories (G) or the latter plus an invariant category (I+G). Amino-acid frequencies were estimated from the input data. The best-fit model according to Bayesian information criterion was JTT+G. Class I and class II monophyly were constrained. 2000 bootstrap replicates were performed with the best model. The resulting phylogeny is available as Supplementary Data 5.

### Giraffe and Okapi repertoire statistics

To isolate OR orthologs lineages, we identified clades in the gene tree with a root node representing the most recent species bipartition (Supplementary Figure 6). Next, the ancestral states of OR genes (namely, whether the ancestral gene was functional one or a pseudogene) were inferred for each node of the tree by assuming that when the node had at least a functional descendant, its state should have been functional. If not, the ancestor node state was labelled as a pseudogene (Supplementary Figure 6). For the subsequent analyses, we subset the orthologous gene lineages to keep only the subtrees with all node bootstrap supports over 90%.

For the histogram representation of Figure 5C, we counted the lineages that have undergone zero (one-to-one orthologs), one, two, three, etc. gene duplications (including pseudogenic paralogs). The species-specific duplication frequency *p(dup)* was calculated for each species as the total number of duplication events normalized by the total number of gene lineages.

To evaluate pseudogenization rates within duplicated genes, we identified all duplication events leading to at least one functional copy (pseudogene duplications were ignored) in each of the orthologous gene lineage. These duplications events were sorted into two categories: fully functional duplication, when two functional copies were produced, and partially functional duplication, when one of the two copies was a pseudogene. For each species, the pseudogenization rate within duplicated genes *p(ps_paralog_)* was calculated as the proportion of partially functional duplications. To calculate the proportion of pseudogenes within one-to-one orthologs *p(ps_ortholog_)*, for each species, we divided the number of pseudogenes by the total number of one-to-one orthologs.

The statistical evaluation of the differences measured between the giraffe and the okapi (i.e. *p(dup)*, *p(ps_paralog_)* and *p(ps_ortholog_)* ratios) consisted in testing if these differences followed the respective amount of molecular divergence accumulated since the two species’ most recent common ancestor, as informed by UCE-calibrated phylogeny.

For pseudogenization frequencies (*p(ps_paralog_)* and *p(ps_ortholog_)*), the probability of the underlying process (genome mutations) was assumed to follow a Poisson process.

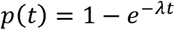

Where *t* represents the time (i.e. the evolutionary distance), *p(t)* the change probability over time (which translates into the observed proportion of pseudogenes) and λ is the change rate. Therefore, the change rate ratio between the two lineages (*Gc* for the giraffe and *Oj* for the okapi) should be equal to:

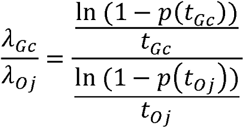

And in the case of equivalent change rates (in this case, pseudogenization rates), we can expect the evolutionary distance ratio to be equal to:

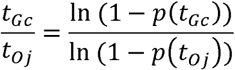

For the duplication rates, we assumed that all duplications could be traced and simply compared the *p(dup)* ratio to the molecular divergence ratio.

To estimate the expected molecular divergence ratio, we divided the corresponding branch lengths of the UCE phylogeny. To test if the duplication or the pseudogenization ratios were significantly different than the expected divergence ratio, we resampled 250 times the orthologous lineages to produce a bootstrap distribution of the ratios. We considered the difference significant if the expected molecular divergence ratio was outside of the 95% interval of the bootstrap distribution.

