## Supplementary Information for "Nose-to-ground distance drives the evolution of mammalian odorant receptor repertoire size"

This document includes supplementary figures 1 to 6 with their corresponding legends, the titles and corresponding legends of supplementary tables 1 to 4 which are grouped in the excel file *Tuberosa_et_al_2025-supplementary_tables.xlsx,* and the titles and description of supplementary data files 1 to 5.

### Data and code availability

All custom scripts, Hidden Markov models, coordinates of functional and pseudogenic coding sequence alignments, and phylogenies generated in this study will be available upon publication.

### Supplementary Figures

#### Supplementary Figure 1

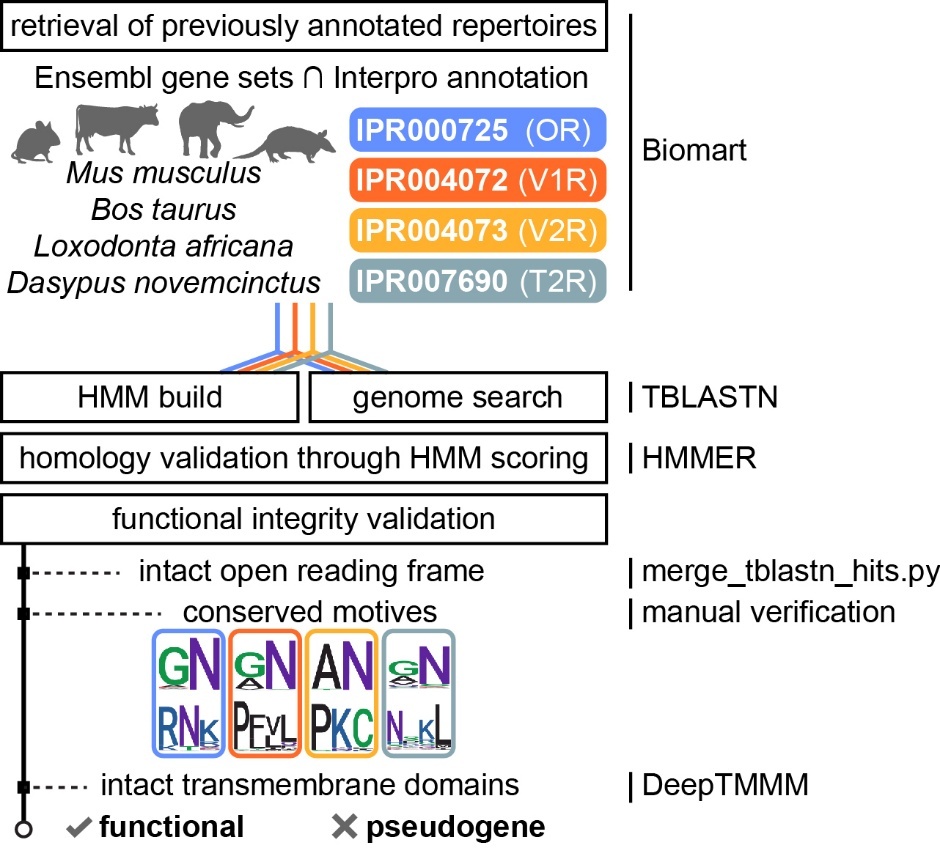

**Pipeline of functional and pseudogenic coding sequence identification.** In summary, model protein sequences of odorant receptors (OR), type 1 vomeronasal receptors (V1R), type 2 vomeronasal receptors (V2R) and bitter taste receptors (T2R) we retrieved from Ensembl using their Interpro annotation. These sets of sequences were used to build HMM models for each receptor family and to search homologous sequences in our genome set via TBLASTN. Homology of the TBLASTN matches was validated using the HMM models, and functionality (whether the sequence should be considered as a pseudogene or not) was inferred by assessing the protein sequence integrity.

#### Supplementary Figure 2

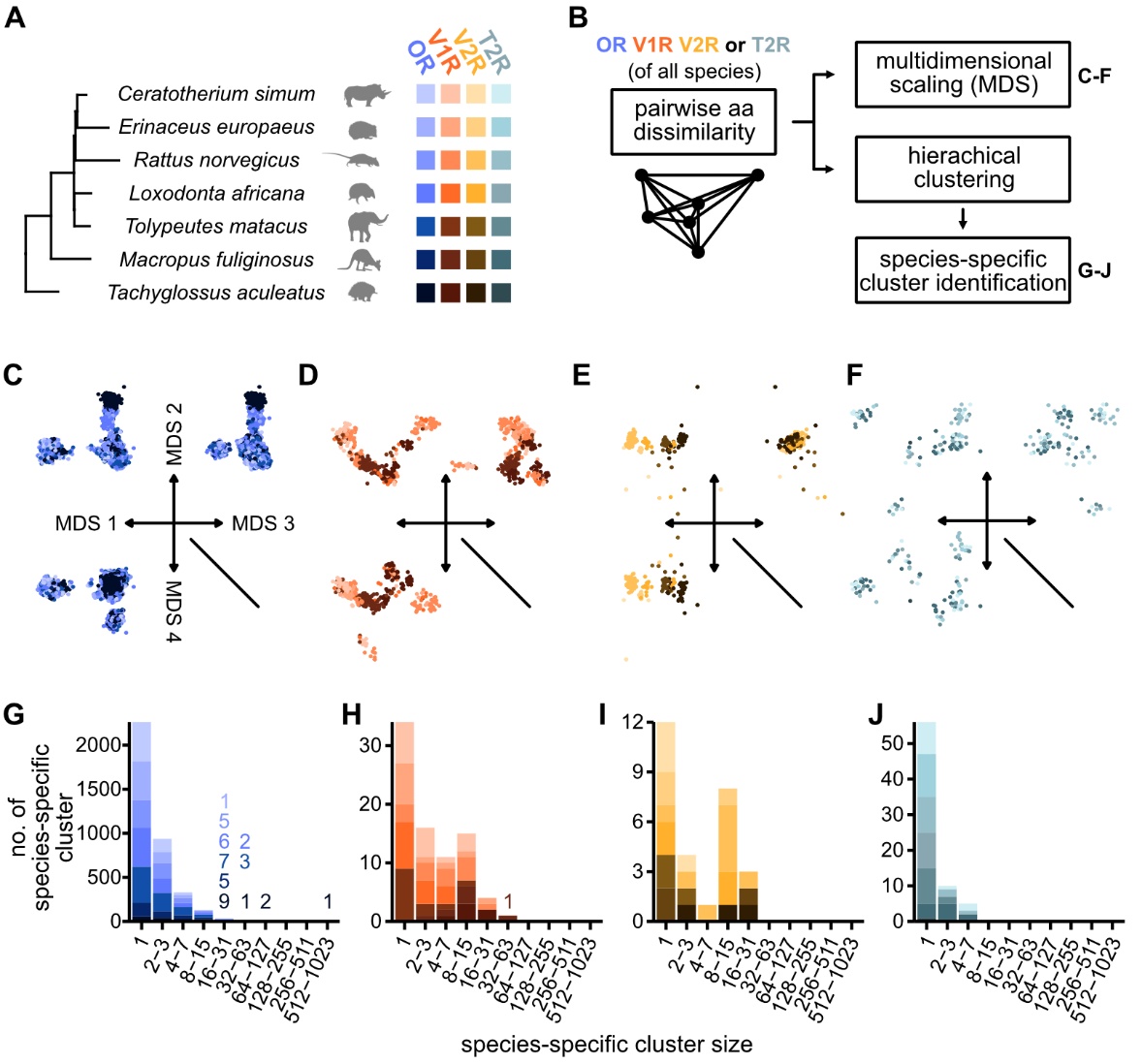

**Evolution dynamics of olfactory and gustatory receptor repertoire identities.** (**A**) The repertoires of functional OR, V1R, V2R and T2R from a subset of seven representative mammal species (the same as in Figure 1B) were used in the subsequent analyses presented in C-J (n=8985 ORs; 396 V1Rs; 200 V2Rs; 106 T2Rs). (**B**) Pairwise amino-acid dissimilarity (see Methods) was calculated for each receptor family, after pooling the seven species repertoires. The resulting dissimilarity matrices were used in two different downstream analyses: a metric multidimensional scaling and a hierarchical clustering. (**C**-**F**) Variance of each of the four dissimilarity matrices was reduced into three orthogonal axes by metric multidimensional scaling (**C**: ORs, **D**: V1Rs, **E**: V2Rs, **F**: T2Rs). (**G**-**J**) Species-specific receptor clusters were identified by hierarchical clustering of the receptor pairwise dissimilarities. The histograms show the distribution of the size of these clusters (namely, the number of receptors they contained) in each receptor family (**G**: ORs, **H**: V1Rs, **I**: V2Rs, **J**: T2Rs). For histogram bars with heights less than 1/50th of the tallest bar, the count values are displayed numerically above their corresponding categories. These numbers maintain the same color shading and stacking arrangement as their respective groups.

#### Supplementary Figure 3

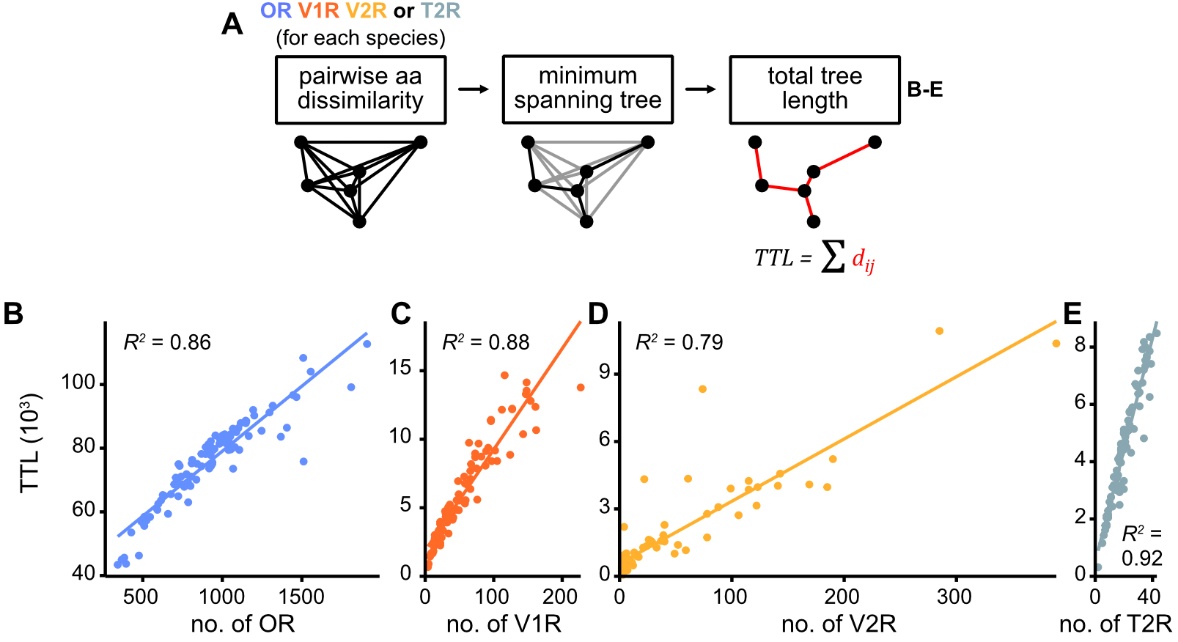

**Chemosensory repertoire diversity and gene numbers.** (**A**) The repertoire of each receptor family and from each species of the main set was analyzed separately, aiming to calculate the total tree length (TTL) of the minimum spanning tree of its pairwise dissimilarities. (**B**-**E**) Linear relationship between the TTL and the number of receptors in the repertoire (**B**: ORs, **C**: V1Rs, **D**: V2Rs, **E**: T2Rs). The adjusted R-squared is shown for each correlation and is strongly significant (p-values < 1e-16).

#### Supplementary Figure 4

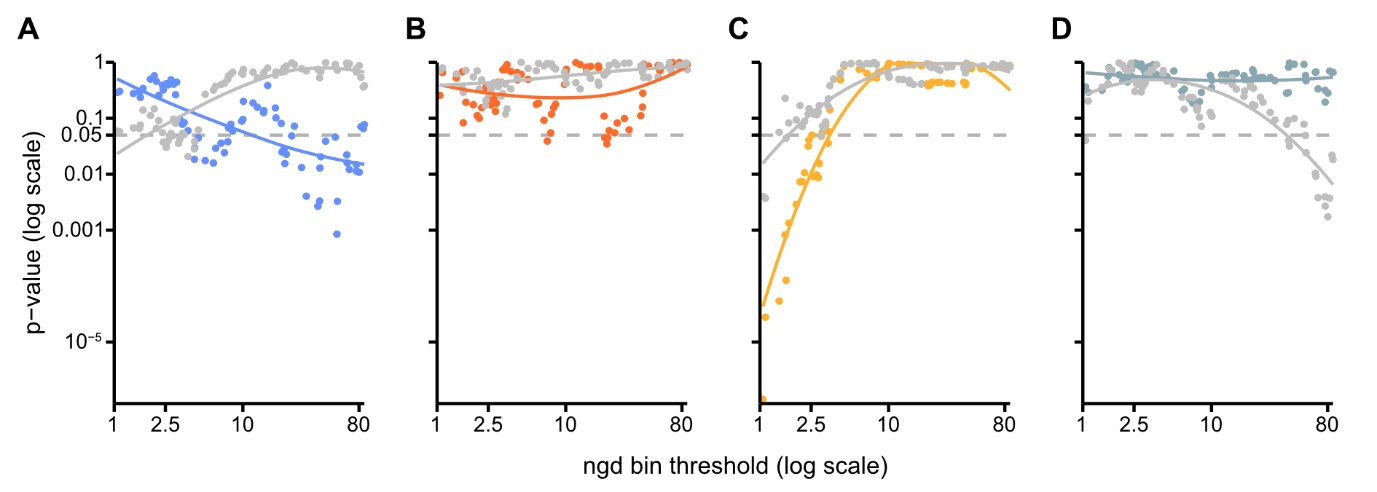

**The number of type 2 vomeronasal receptors maybe influenced by the nose-to-ground distance.** (**A**-**D**) P-value (y-axis) of a pGLS regression comparing the number of functional genes between two categories of nose-to-ground distance, close and far, defined by a threshold value (x-axis). (**A**) ORs. (**B**) V1Rs. (**C**) V2Rs. (**D**) T2Rs.

#### Supplementary Figure 5

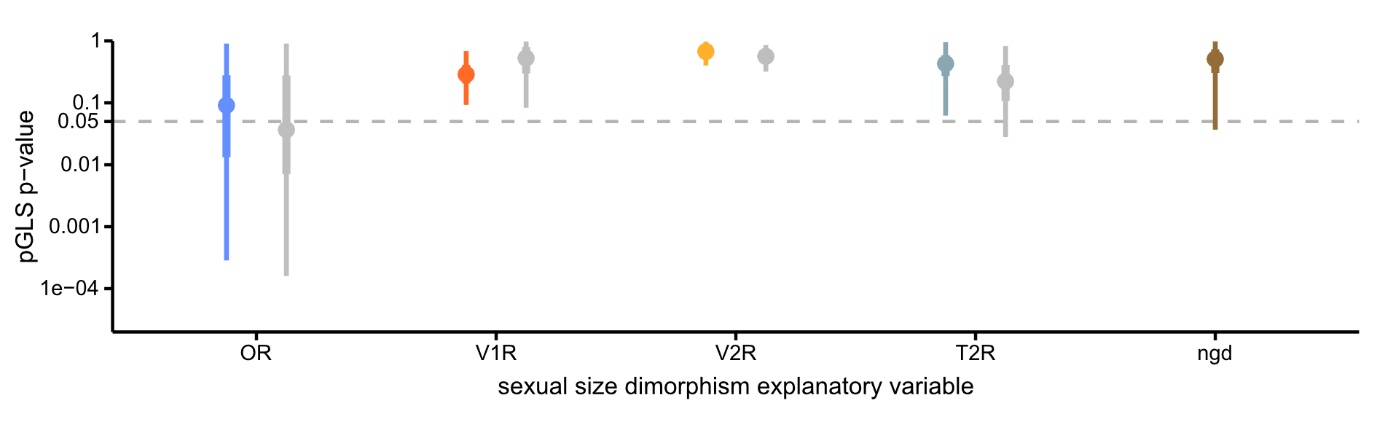

**Sexual size dimorphism may predict the number of functional OR and OR pseudogenes.** P-value ranges of the pGLS regression of the number of gene by the sexual-size difference in alternative species sets. Mean n=28.1 (standard deviation=1.7).

**Supplementary Figure 6**

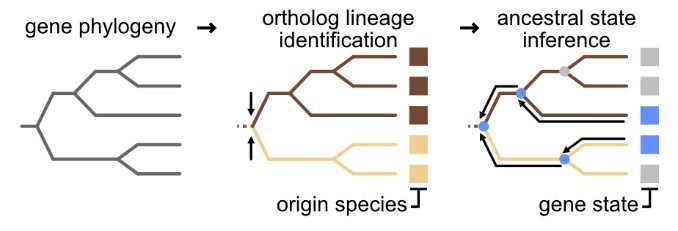

**Giraffe and okapi OR gene ancestral state reconstruction.** First, the gene phylogeny is obtained from the multiple alignment of amino acids sequences from OR functional and pseudogenic CDSs, using an ML approach. Next, the origin of all orthologous gene lineages in the tree are identified by finding all nodes joining two different species-specific clades. Finally, the ancestral states of OR genes (namely, whether the ancestral gene was functional one or a pseudogene), is inferred by assuming a simple transition model, independent of branch lengths, where functional state was attributed when there was at least a functional descendant. If not, the ancestor node state is labelled as a pseudogene.

### Supplementary Tables

**Supplementary Table 1**

**List of analyzed species** with their respective NCBI’s Taxonomy identifiers, family and phylogenetic grouping (used in boxplot and linear regression displays). The three last columns identify species featured in the main 96 species set, in the MDS and hierarchical clustering analyses of receptor identities, and with their OR genomic proportions in Figure 1B, respectively.

#### Supplementary Table 2

**List of analysed genome assemblies**, with their Genbank or RefSeq identifier, and their BUSCO analysis values.

**Supplementary Table 3**

**Animal traits identified for this study**, and their corresponding exclusive categories into which species were sorted.

#### Supplementary Table 4

**Species metadata**, including nose-to-ground distance (ngd), activity period, diet, territoriality, sociality and numbers of functional or pseudogenic receptors.

### Supplementary Data

#### Supplementary Data 1

Species pictures with nose-to-ground distance measurements.

Supplementary Data 1 will be available upon publication.

#### Supplementary Data 2

UCE-calibrated species phylogeny, full species set (n spp=158).

#### Supplementary Data 3

UCE-calibrated species phylogenies, 250 alternative species sets (n spp=96).

#### Supplementary Data 4

Coordinates of functional and pseudogenic receptor coding sequences.

#### Supplementary Data 5

Giraffe and okapi’s OR phylogeny.
